# The single-cell transcriptional landscape of lung carcinoid tumors

**DOI:** 10.1101/2021.12.07.471416

**Authors:** Philip Bischoff, Alexandra Trinks, Jennifer Wiederspahn, Benedikt Obermayer, Jan Patrick Pett, Philipp Jurmeister, Aron Elsner, Tomasz Dziodzio, Jens-Carsten Rückert, Jens Neudecker, Christine Falk, Dieter Beule, Christine Sers, Markus Morkel, David Horst, Frederick Klauschen, Nils Blüthgen

## Abstract

Lung carcinoid tumors, also referred to as pulmonary neuroendocrine tumors or lung carcinoids, are rare neoplasms of the lung with a more favorable prognosis than other subtypes of lung cancer. Still, some patients suffer from relapsed disease and metastatic spread while no consensus treatment exists for metastasized carcinoids. Several recent single-cell studies have provided detailed insights into the cellular heterogeneity of more common lung cancers, such as adeno- and squamous cell carcinoma. However, the characteristics of lung carcinoids on the single-cell level are yet completely unknown.

To study the cellular composition and single-cell gene expression profiles in lung carcinoids, we applied single-cell RNA sequencing to three lung carcinoid tumor samples and normal lung tissue. The single-cell transcriptomes of carcinoid tumor cells reflected intertumoral heterogeneity associated with clinicopathological features, such as tumor necrosis and proliferation index. The immune microenvironment was specifically enriched in noninflammatory monocyte-derived myeloid cells. Tumor-associated endothelial cells were characterized by distinct gene expression profiles. A spectrum of vascular smooth muscle cells and pericytes predominated the stromal microenvironment. We found a small proportion of myofibroblasts exhibiting features reminiscent of cancer-associated fibroblasts. Stromal and immune cells exhibited potential paracrine interactions which may shape the microenvironment via NOTCH, VEGF, TGFβ and JAK/STAT signaling. Moreover, single-cell gene signatures of pericytes and myofibroblasts demonstrated prognostic value in bulk gene expression data.

Here, we provide first comprehensive insights into the cellular composition and single-cell gene expression profiles in lung carcinoids, demonstrating the non-inflammatory and vessel-rich nature of their tumor microenvironment, and outlining relevant intercellular interactions which could serve as future therapeutic targets.

## Introduction

Lung cancer is a heterogeneous disease comprising different histopathological subtypes. Besides adenocarcinomas and squamous cell carcinomas, the 2015 WHO classification established the category of pulmonary neuroendocrine tumors (NETs) [1]. This category comprises the high-grade subtypes small cell lung cancer (SCLC) and large cell neuroendocrine carcinoma (LCNEC), and the low- and intermediate-grade NETs of the lung, also referred to as typical and atypical carcinoids, respectively. Lung carcinoids contribute to 1 % of lung cancer cases [2] with an increasing incidence over the last decades [3]. On average, lung carcinoids have a better outcome than conventional lung cancers. Typical carcinoids and atypical carcinoids, of which the latter are specified by higher mitotic rate or presence of tumor necrosis, have a 5-year survival rate of approximately 90% and 70%, respectively [3, 4]. About 10% of carcinoid patients present with regional lymph node metastasis [3, 5]. Atypical carcinoids have a higher risk of lymphonodal and systemic metastatic spread, and recurrent disease [5, 6]. However, no consensus exists for a standardized systemic therapeutic regimen of metastasized lung carcinoids [7].

The more common subtypes of lung cancer, i.e., adenocarcinomas and squamous cell carcinomas, are related to smoking and characterized by high tumor mutational burden. In contrast, lung carcinoids affect younger patients and non-smokers, harbor a significantly lower mutational load and a different spectrum of oncogenic mutations [8]. Consequently, novel targeted and immune therapies, which have already improved the outcome in lung adeno- and squamous cell carcinomas [9], cannot easily be translated to lung carcinoids. Moreover, predicting the efficacy of modern targeted and immune therapies is limited by intratumoral heterogeneity, where tumors may harbor primary resistant tumor cell subclones, as well as the complex tumor microenvironment, modulating immune responses against the tumor. Single-cell gene expression profiling allows to overcome this limitation and has already provided valuable insights into the cellular heterogeneity of lung adenocarcinomas [10–14].

In this study, we comprehensively analyzed the cellular composition of lung carcinoids by applying single-cell RNA sequencing to three carcinoid tumor and normal lung tissue samples. We show that single-cell gene expression profiles of carcinoid tumor cells reflect clinicopathological features and allow assignment to recently defined molecular clusters [15]. Further, we found that the tumor microenvironment was characterized by differentiating monocyte-derived myeloid cells with non-inflammatory features, tumor-associated endothelial cells, a spectrum of vascular smooth muscle cells and pericytes, and myofibroblasts with cancer-associated fibroblast-like features. Our analysis provides the basis for further studies of the lung carcinoid tumor microenvironment, potential prognostic and predictive biomarkers as well as novel therapeutic targets.

## Results

### Single-cell RNA sequencing uncovers the cellular diversity of lung carcinoids

To explore the cellular composition of lung carcinoids and their tumor microenvironment on the single-cell level, we collected fresh tissue samples of tumor tissue and normal lung parenchyma from 3 previously untreated lung carcinoid patients undergoing primary surgery (patients 1-3, Fig. 1A). All three patients showed tumor cells growing in solid nests with expression of the neuroendocrine marker proteins synaptophysin and chromogranin A (Fig. 1B). Tumors comprised one typical carcinoid (patient 1), one atypical carcinoid with high proliferative activity (patient 2, see Supp. Fig. 1A for Ki67 immunostaining), and one atypical carcinoid with focal tumor necrosis (patient 3, see Supp. Fig. 1B for HE staining of necrotic area). Both atypical carcinoid cases (patients 2 and 3) had regional lymph node metastases at the time of diagnosis. Tissue samples were enzymatically dissociated and subjected to singlecell RNA sequencing using a commercial droplet-based system. Single-cell gene expression data of 7 normal lung tissue samples from a previously published cohort (patients 4-10) [14] were included in the subsequent analyses. Altogether, we analyzed 73,105 single-cell transcriptomes of which 64,697 high-quality transcriptomes remained after quality control and filtering (Fig. 1C, see Supp. Fig. 1C-D for quality control parameters).

**Figure 1:**
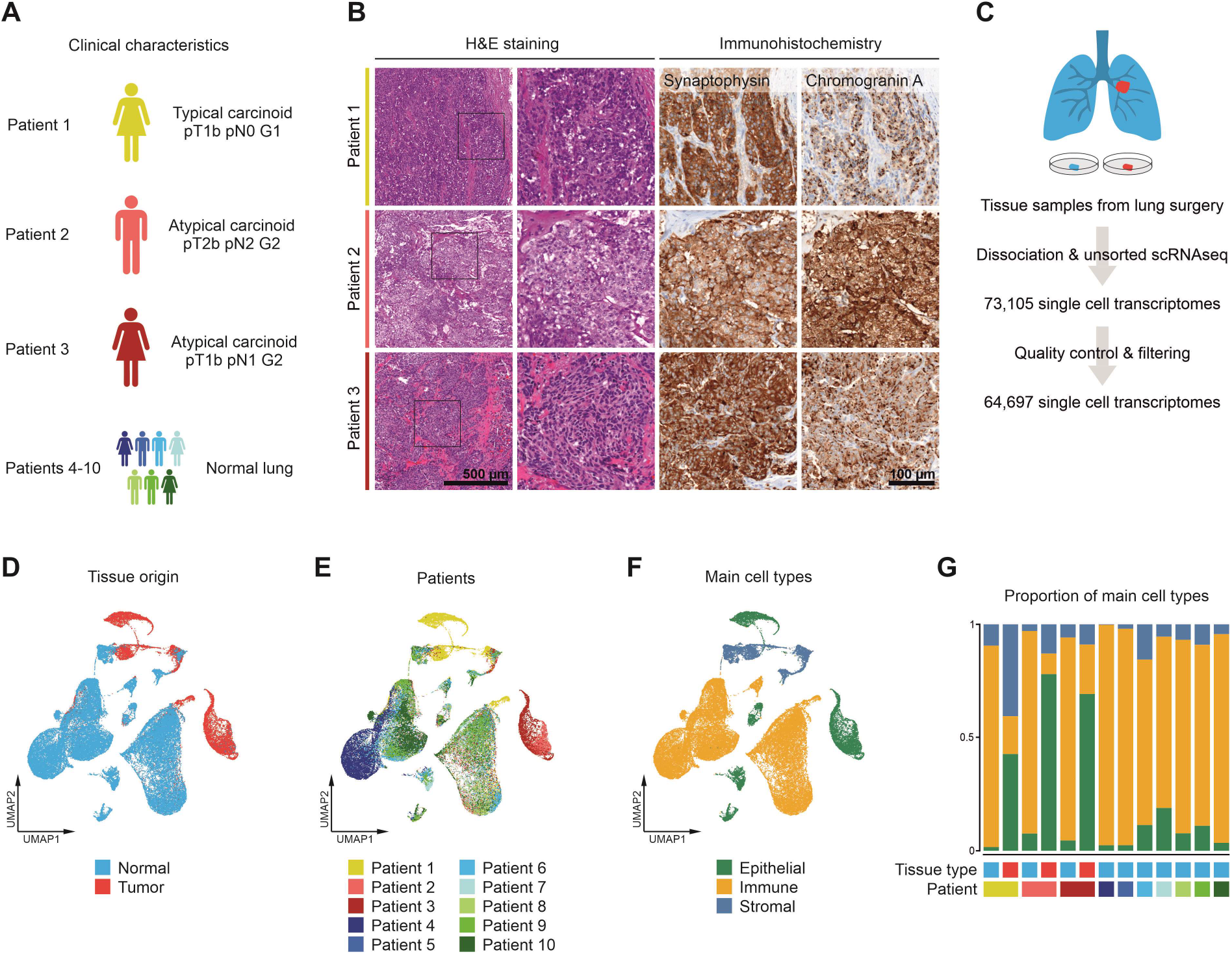
Single-cell RNA sequencing of lung carcinoids. (A) Clinical characteristics of 3 lung carcinoid patients analyzed in this study. (B) H&E staining and immunohistochemical staining for neuroendocrine marker proteins of 3 lung carcinoid patients. (C) Simplified schematic representation of the single-cell RNA sequencing workflow. (D-F) UMAPs based on the top 10 principal components of all single-cell transcriptomes after filtering, color-coded by (D) tissue type, (E) patients, and (F) main cell type. (G) Proportions of main cell types per sample.

Visualization of single-cell transcriptomes by uniform manifold approximation and projection (UMAP) revealed distinct shifts between normal and tumor tissue samples (Fig. 1D). Note that single-cell transcriptomes of different patients overlapped in many clusters, excluding systematic batch effects across samples (Fig. 1E). In the epithelial, immune and stromal cell compartment, which were defined by gene expression of canonical marker genes (Supp. Fig. 1E), we observed tumor-specific changes (Fig. 1F). In the tumor tissue samples, we mostly found epithelial and stromal single-cell transcriptomes, whereas immune single-cell transcriptomes were more abundant in the normal tissue samples (Fig. 1G).

### Intertumoral heterogeneity of lung carcinoids reflects clinicopathological features and molecular subtypes

To further analyze the epithelial cell compartment, epithelial single-cell transcriptomes were subset and re-clustered. Epithelial cell clusters overrepresented in normal or tumor tissue samples were assigned as normal or tumor cell clusters, respectively (Supp. Fig. 2A, 2B). We observed that normal cell clusters were shared by different patients whereas tumor cell clusters were highly patient-specific (Fig. 2A). In the normal cell clusters, using canonical marker genes and predefined gene signatures [16, 17], we identified alveolar epithelial type 1 and 2, ciliated, club, and basal cells (Fig. 2A-B, Supp. Fig. 3A-B). As indicated by the highly patient-specific tumor cell clusters, we found many differentially expressed genes in the tumor cells between patients (Fig. 2C), such as *CD44*, *TFF3* and *EGFR*, which correlated with differential protein expression, as shown by immunohistochemistry (Fig. 2D). Highly expressed genes in the typical carcinoid of patient 1 comprised many that have been associated with good prognosis, such as *MT1G, MT1M, MT1X, PCK1, LPL, CD44* [18]. While we found distinct transcriptional differences between different tumor cases, transcriptional profiles within individual tumors were quite homogeneous and varied mainly depending on the number of reads and genes per cell (Supp. Fig. 2C-F).

**Figure 2:**
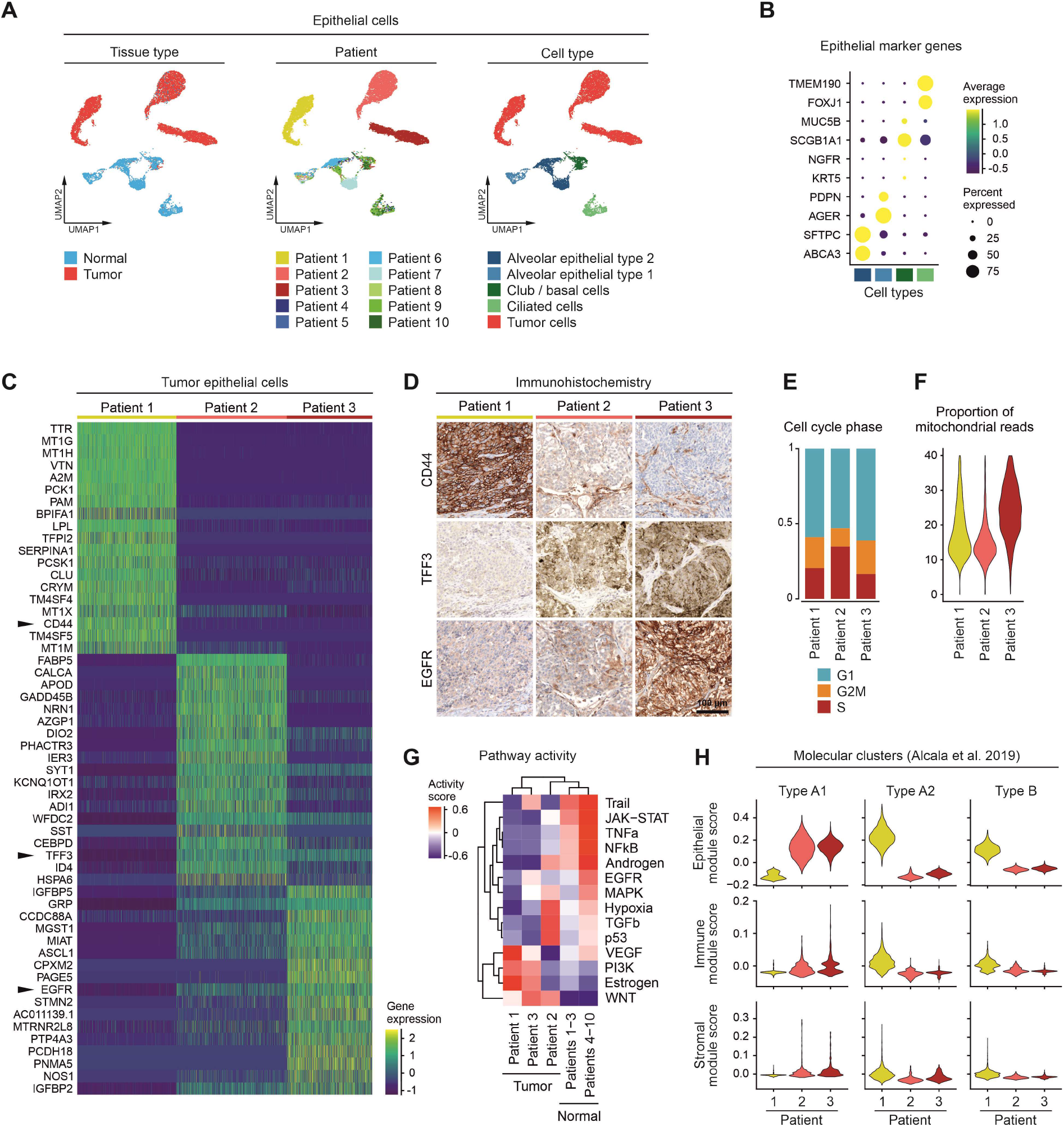
Intertumoral heterogeneity of tumor cells in lung carcinoids. (A) UMAPs based on the top 10 principal components of all epithelial single-cell transcriptomes color-coded by tissue type, patient, and cell type. (B) Average gene expression of selected marker genes of normal epithelial cell types, for cell type color code see (A). (C) Differentially expressed genes in tumor epithelial cells grouped by patients, top 20 genes showed per patient, for patient color code see (A). (D) Immunohistochemical staining of proteins encoded by selected differentially expressed genes indicated by black arrowheads in (C). (E) Proportion of tumor epithelial cells assigned to different cell cycles, grouped by patient. (F) Proportion of mitochondrial reads in tumor epithelial transcriptomes, grouped by patient. (G) Mean pathway activity scores of tumor epithelial cells, grouped by patient, and normal epithelial cells, grouped by patient groups. (H) Module scores of marker genes of molecular clusters according to Alcala et al. [15] in epithelial, immune, and stromal cells, grouped by patient.

To further explore interpatient heterogeneity, we inferred different functional traits from the single-cell gene expression profiles, namely cell cycle phase, proportion of mitochondrial reads and signaling pathway activity. The tumor cells of patient 2 had the highest proportion of cells in S phase while at the same time showing the highest Ki67 proliferation index in immunohistochemistry (Fig. 2E, see Supp. Fig. 1A for Ki67 immunostaining). The highest proportion of mitochondrial reads was observed in tumor cells of patient 3 which was characterized by focal tumor necrosis (Fig. 2F, see Supp. Fig. 1B for H&E of necrotic area). Tumor cell transcriptomes of patient 3 had high scores for EGFR pathway activity and strong EGFR expression on the protein level (Fig. 2D, 2G). The pathway activity scores for estrogen and androgen receptor signaling correlated with the patient’s sex (Fig. 2G, see Fig. 1A for clinical characteristics). Recently, it has been shown that lung carcinoids can be subtyped into three distinct molecular clusters based on transcriptional and epigenetic features [15]. In our single-cell gene expression profiles, we could assign the tumors of patients 2 and 3 to cluster A1, and the tumor of patient 1 both to cluster A2 and cluster B (Fig. 2H). Notably, the immune and stromal cell compartment exhibited only minor expression scores of molecular cluster gene signatures.

Taken together, lung carcinoid tumor single-cell transcriptomes revealed intertumoral heterogeneity which reflected different clinical and histomorphological features, such as patient’s sex, tumor proliferative activity and tumor necrosis, as well as recently proposed molecular clusters of lung carcinoids.

### The immune microenvironment of lung carcinoids is characterized by non-inflammatory monocyte-derived myeloid cells

To discover the cellular composition of the immune microenvironment, immune single-cell transcriptomes were subset and re-clustered. We identified a variety of different cell types within the immune cell compartment using canonical marker genes and predefined gene signatures [16, 17] (Fig. 3A-B, Supp. Fig. 3A-B).

**Figure 3:**
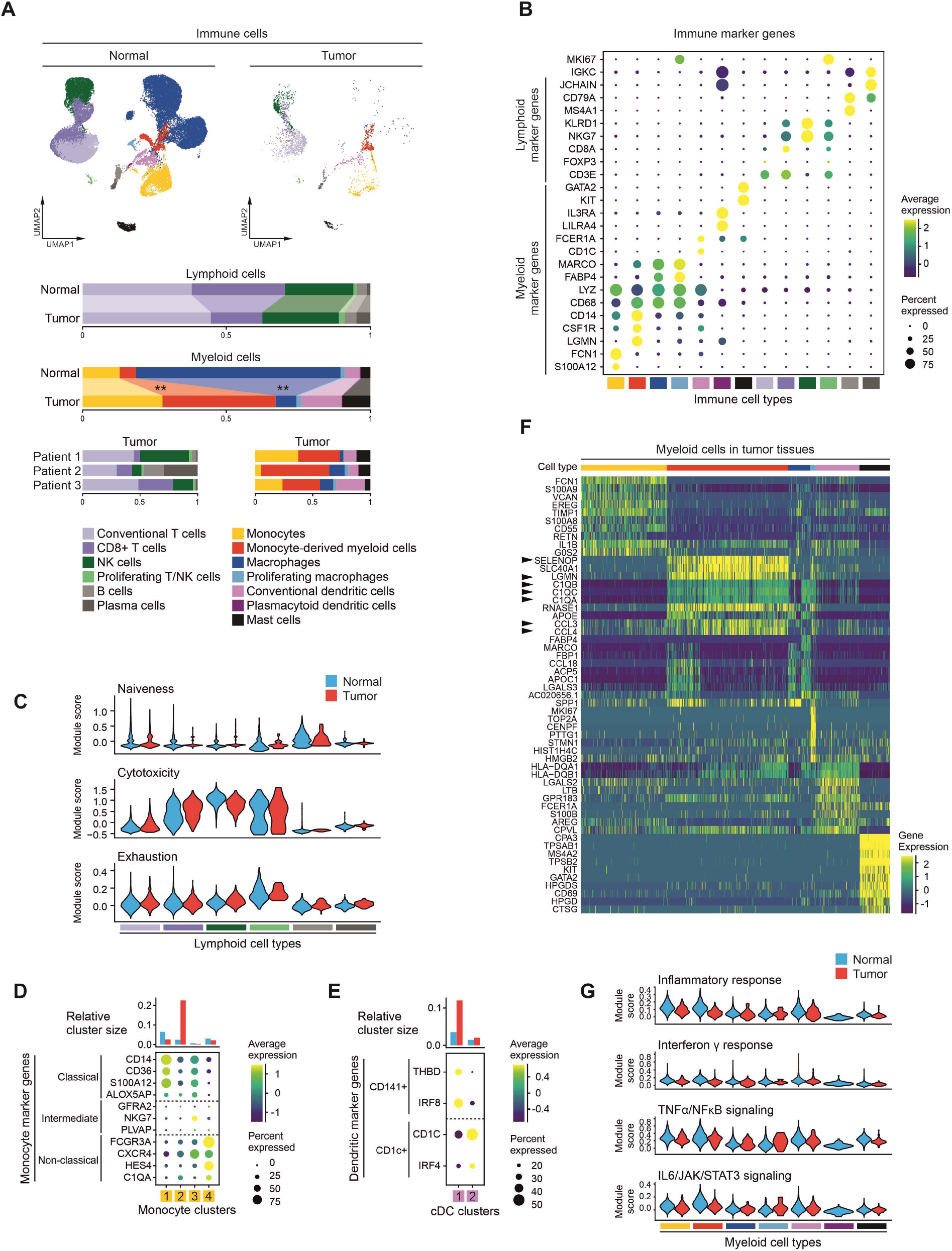
Composition of the immune tumor microenvironment in lung carcinoids. (A) UMAPs based on the top 10 principal components of all immune single-cell transcriptomes, split by tissue type, color-coded by cell type, and proportions of lymphoid and myeloid cell types per tissue type and, for tumor samples, per patient, Mann-Whitney U test, ** = p<0.01. (B) Average gene expression of selected marker genes of immune cell types, for cell type color code see (A). (C) Module scores of gene signatures related to naiveness, cytotoxicity and exhaustion in different lymphoid cell types, split by tissue type, for cell type color code see (A). (D) Average gene expression of selected marker genes of monocyte subsets and relative size of monocyte clusters, for tissue type color code see (C). (E) Average gene expression of selected marker genes of conventional dendritic cell subsets and relative size of conventional dendritic cell clusters, for tissue type color code see (C). (F) Differentially expressed genes in myeloid cells in tumor samples, grouped by cell type, top 10 genes shown per cell type, for cell type color code see (A), black arrowheads indicate genes mentioned in the main text. (G) Module scores of gene signatures related to immune response in different myeloid cell types, split by tissue type, for cell type color code see (A), for tissue type color code see (C).

We identified different lymphoid cells, such as conventional T cells, CD8+ T cells, NK cells, B cells and plasma cells (Fig. 3A, Supp. Fig. 4A-B, Supp. Table 1-2). On average, the proportion of lymphoid cell types in tumor tissues closely resembled normal tissues while we also noted some interpatient heterogeneity in both tumor and normal tissues. Gene signatures reflecting naiveness, cytotoxicity or exhaustion [13] of lymphoid cells showed similar expression scores in normal and tumor tissues (Fig. 3C). Note that signature scores for exhaustion were low in all lymphoid cell types, contrasting the microenvironment of lung adenocarcinoma in a reference dataset, which was enriched in exhausted T cells (Supp. Fig. 5A-B) [14]. Together, the lymphoid microenvironment of lung carcinoids resembles the lymphoid cell compartment of normal lung parenchyma.

Within the myeloid cell compartment, we identified monocytes, dendritic cells, macrophages and mast cells with different abundancies in tumor and normal tissues (Fig. 3A, Supp. Fig. 4A, Supp. Table 3-4). Among the monocytes, the classical monocyte cluster 2 was enriched in tumor tissues (Fig. 3D). Conventional dendritic cells comprised two clusters of which the CD141+ cluster 1 was mostly found in tumor tissues (Fig. 3E). Beyond, we identified a tumor-enriched cell cluster with high expression levels of both monocyte markers, such as *CD14*, and *LGMN*, a gene upregulated in differentiating monocytes [19] (Fig. 3B). We conclude that this cell cluster represents the spectrum of monocyte-derived myeloid cells differentiating either into macrophages, as shown by high *APOC1* and *APOE* expression in cluster 3, or into dendritic cells, as shown by high *S100A8* and *S100A9* expression in cluster 1 [20, 21] (Supp. Fig. 4D). While the proportions of monocytes and conventional dendritic cells were heterogeneously increased across patients, monocyte-derived myeloid cells were consistently increased across all three carcinoid tumors analyzed (p = 0.0070). Monocyte-derived myeloid cells were further characterized by high expression of *SELENOP, C1QA, C1QB, C1QC* and the chemokines *CCL3* and *CCL4* (Fig. 3F). Compared to normal tissues, monocyte-derived myeloid cells in tumor tissues showed equal to slightly lower expression scores of various gene signatures related to inflammation and immune response (Fig. 3G). In a reference dataset of lung adenocarcinoma, the microenvironment was composed of both pro- and non-inflammatory clusters (Supp. Fig. 5A-B). In contrast, our results indicate that the lung carcinoid immune microenvironment is predominated by non-inflammatory monocyte-derived myeloid cells.

The composition of the myeloid cell compartment was to some degree heterogeneous across the three lung carcinoids analyzed by single-cell RNA sequencing. In order to study interpatient heterogeneity in a larger cohort, we quantified the expression of marker genes of characteristic immune cell types in a published bulk gene expression dataset of lung carcinoids [15, 22]. Here, we found that marker genes of the tumor-enriched monocyte-derived myeloid cell cluster 2 and 3 were associated with atypical carcinoids, albeit not correlated with overall survival (Supp. Fig. 4E), indicating microenvironmental differences between lung carcinoid subtypes.

### Vascular cells and CAF-like myofibroblasts constitute the stromal microenvironment of lung carcinoids

To gain insight into the composition of the stromal microenvironment, stromal single-cell transcriptomes were subset and re-clustered. Here, we identified different clusters of endothelial, fibroblastic and smooth muscle cells using canonical marker genes and predefined gene signatures [16, 17] (Fig. 4A–4B, Supp. Fig. 3A-B).

**Figure 4:**
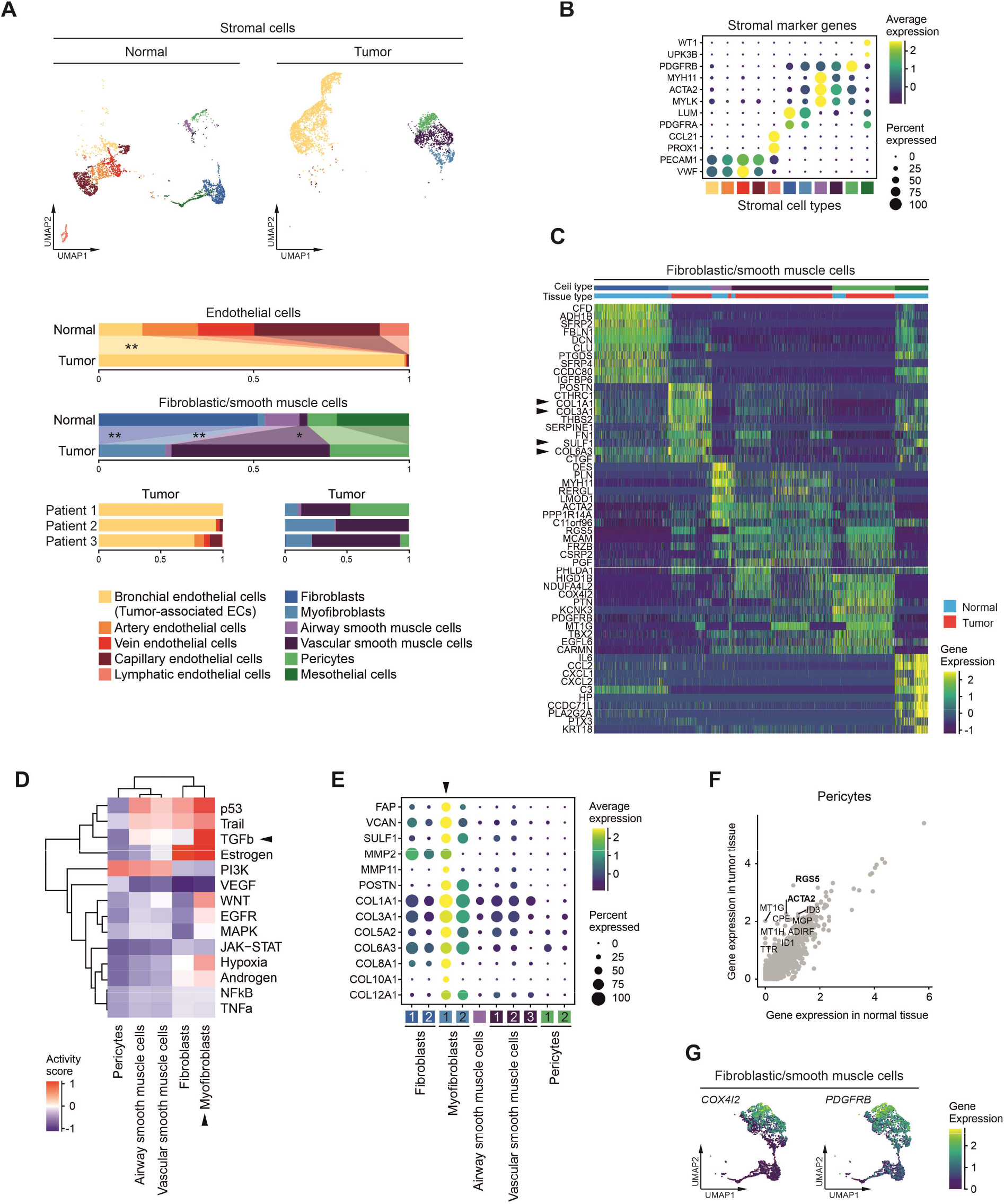
Composition of the stromal tumor microenvironment in lung carcinoids. (A) UMAPs based on the top 10 principal components of all stromal single-cell transcriptomes split by tissue type, color-coded by cell type, and proportions of endothelial and fibroblastic/smooth muscle cell types per tissue type and, for tumor samples, per patient, Mann-Whitney U test, * = p<0.05, ** = p<0.01. (B) Average gene expression of selected marker genes of stromal cell types, for cell type color code see (A). (C) Differentially expressed genes in fibroblastic/smooth muscle cells, grouped by cell type, top 10 genes shown per cell type, for cell type color code see (A), black arrowheads indicate genes mentioned in the main text. (D) Mean pathway activity scores of different fibroblastic/smooth muscle cell clusters, mesothelial cells excluded, black arrowheads indicate pathways and cell types mentioned in the main text. (E) Average expression of selected marker genes of myofibroblast cluster 1 as indicated by black arrowhead. (F) Average gene expression of pericytes in tumor versus normal tissues, top 10 genes overexpressed in tumor tissues indicated, genes mentioned in the main text in bold. (G) UMAPs of fibroblastic/smooth muscle cells, colored by gene expression of canonical pericyte marker genes.

Among the endothelial cells we could distinguish bronchial, capillary, arterial and venous endothelial cells based on predefined marker gene signatures [16] (Fig. 4A, Supp. Fig. 3A, Supp. Table 5-6). While different subtypes of endothelial cells were present in normal lung parenchyma, tumor tissues were significantly enriched in bronchial-type endothelial cells (p = 0.0070) (Fig. 4A, Supp. Fig. 6A). Here, the majority of endothelial transcriptomes was obtained from the tumor of patient 1, correlating with dense vascularization as shown by immunostaining (Supp. Fig. 6B). Endothelial cells in tumor tissues showed high mRNA expression of *INSR*, a marker gene of tumor-associated endothelial cells [23], and high INSR protein expression, contrasting normal lung tissue (Supp. Fig. 6C-D). Moreover, we found high expression of genes that have been related to angiogenesis, such as *VWA1*, *COL15A1*, *IGFBP7* and *GSN* [10] (Supp. Fig. 6D), suggesting a phenotype of tumor-associated endothelial cells in the microenvironment of lung carcinoids comparable to those found in the microenvironment of lung adenocarcinoma (Supp. Fig. 5A, C) [14].

Within the fibroblastic and smooth muscle cell compartment in tumor tissues, we found myofibroblasts, vascular smooth muscle cells and pericytes, whereas fibroblasts were significantly decreased compared to normal tissues (p = 0.0091) (Fig. 4A, Supp. Table 7-8). Myofibroblasts were strongly enriched in tumor tissues (p = 0.0074) and showed high expression of extracellular matrix components, such as *COL1A1*, *COL3A1* and *COL6A3*, as well as matrix-degrading enzymes, such as *SULF1* (Fig. 4C), suggesting that these cells might be involved in extracellular matrix remodeling. Moreover, myofibroblasts were characterized by high activity of TGFβ signaling (Fig. 4D). Within the myofibroblasts, cluster 1 showed a specific overexpression of *FAP* and *MMP11* as well as a higher expression of various collagens, compared to myofibroblast cluster 2 and other fibroblast and smooth muscle cell clusters (Fig. 4E). In an independent cohort of lung carcinoids characterized by bulk RNA sequencing [15, 22], the marker gene signature of myofibroblast cluster 1 was significantly associated with atypical carcinoids and correlated with worse overall survival (Supp. Fig. 6E). Together, these data indicate that myofibroblasts in lung carcinoid tumor tissues exhibit biological traits characteristic of cancer-associated fibroblasts.

The stromal microenvironment of carcinoids was predominated by vascular smooth muscle cells (p = 0.0160) and pericytes, contrasting the microenvironment of lung adenocarcinoma which mainly contained myofibroblasts (Supp. Fig. 5A, C) [14]. The highest proportion of pericytes was found in case 1 which was diagnosed as a typical carcinoid whereas fewer pericytes were found in patients 2 and 3 diagnosed with atypical carcinoids (Fig. 4A). Correspondingly, in an independent lung carcinoid cohort characterized by bulk RNA sequencing [15, 22], the pericyte marker gene signature was associated with typical carcinoids and correlated with better overall survival (Supp. Fig. 6F). Pericytes in tumor tissues showed a high expression of *RGS5*, a gene involved in pericyte development, and *ACTA2*, a smooth muscle marker gene (Fig. 4F). Smooth muscle cells expressed low levels of pericyte marker genes, such as *COX4I2* and *PDGRB* (Fig. 4C). We found these genes expressed in a graded fashion suggesting that pericytes and vascular smooth muscle cells rather form a continuum than discrete cell types in tumor tissues (Fig. 4G). These results show that the stromal microenvironment of lung carcinoids is composed of myofibroblasts reminiscent of cancer-associated fibroblasts, and a spectrum of vascular smooth muscle cells and pericytes. Myofibroblasts and pericytes may be linked to worse and better overall survival, respectively.

### Interactions between tumor microenvironmental cells potentially activate NOTCH, VEGF, TGFβ, and JAK/STAT signaling

We observed that different cell types are specifically enriched in the lung carcinoid microenvironment (Fig. 5A). In order to delineate functional relationships between microenvironmental and tumor cells, we quantified potential paracrine receptor-ligand interactions [24]. Interestingly, most potential interactions were found between cell types of the stromal microenvironment, involving tumor-associated endothelial cells, myofibroblasts, vascular smooth muscle cells and pericytes, whereas tumor cells are less involved in potential paracrine interactions (Fig. 5B, Supp. Table 9). Note that the number of potential interactions was independent from the number of cells or mean number of mRNA counts per cell type (Supp. Fig. 7A-B). Focusing on the most relevant signaling pathways, we found many interactions potentially activating NOTCH, TGFβ, VEGF, and JAK/STAT signaling (Fig. 5C). Tumor-associated endothelial cells receive potentially VEGF, TGFβ, and NOTCH pathway-activating signals, while myofibroblast mainly receive potentially TGFβ pathway-activating signals, both via various paracrine and autocrine interactions. Dendritic cells, monocytes and monocyte-derived myeloid cells receive potentially JAK/STAT pathway-activating signals mainly via autocrine and paracrine interactions with other immune cells. These results indicate that autocrine and paracrine interactions between various stromal and immune cells may shape the lung carcinoid tumor microenvironment.

**Figure 5:**
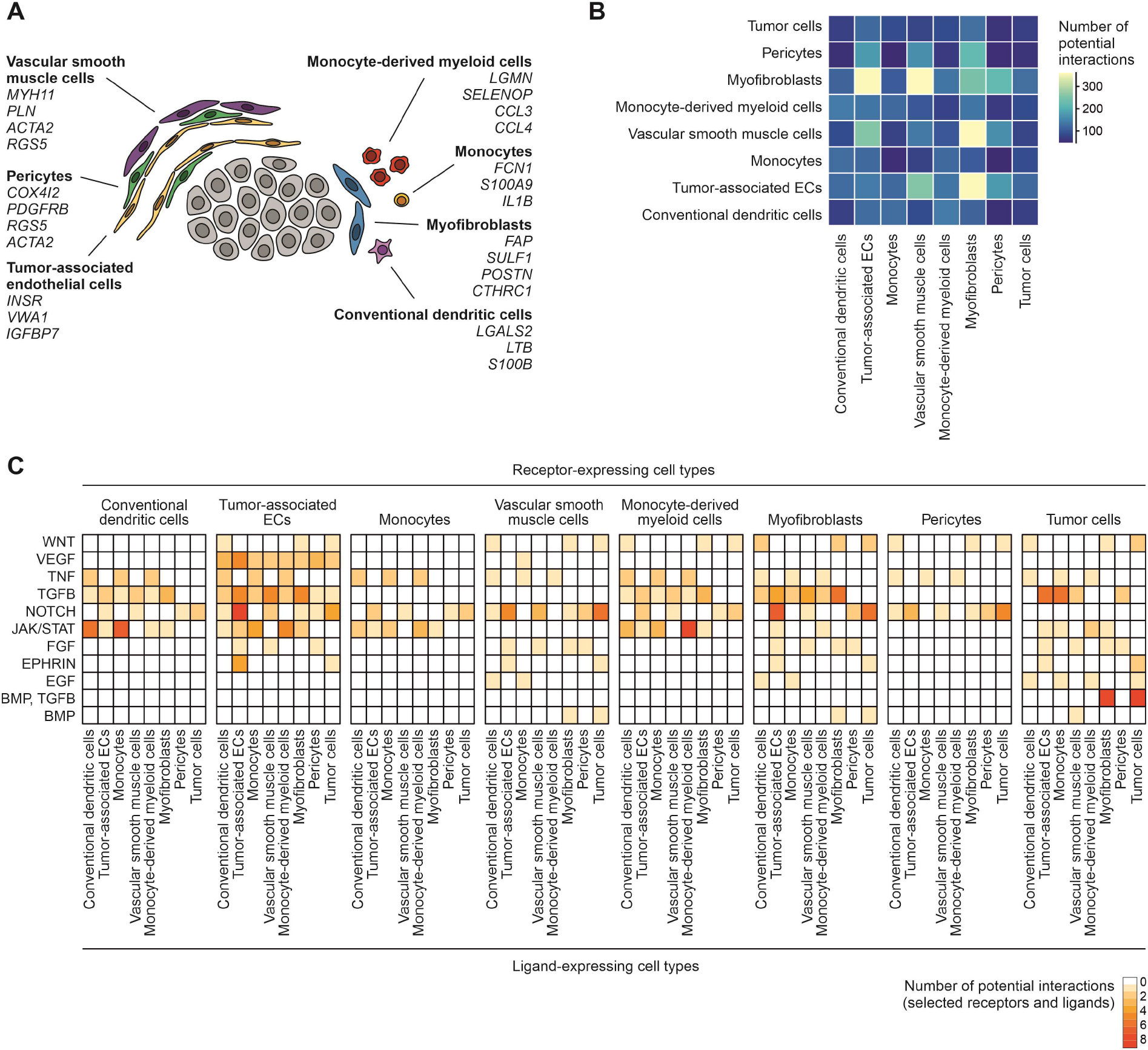
Potential paracrine interactions within the lung carcinoid tumor microenvironment. (A) Characteristic cell types of the lung carcinoid tumor microenvironment and selected cell type marker genes. (B) Number of potential auto-/paracrine interactions between characteristic cell types of the lung carcinoid tumor microenvironment, calculated using the CellPhoneDB algorithm. (C) Number of potential cell-cell interactions filtered for high-confidence receptors and ligands of relevant signaling pathways, grouped by interaction families. Each heatmap shows potential interactions where the respective receptor is expressed in the cell type indicated above.

## Discussion

In this study, we analyzed the tumor heterogeneity and cellular composition of the tumor microenvironment in lung carcinoids. By applying single-cell RNA sequencing to tumors from three patients, we outline the single-cell landscape of lung carcinoids in unprecedented depth and comprehensiveness. We could show that tumor cell transcriptomes reflect high intertumoral but low intratumoral heterogeneity. The immune microenvironment was characterized by non-inflammatory monocyte-derived myeloid cells, classical monocytes and conventional dendritic cells, while the lymphoid cell compartment was comparable to normal lung parenchyma. The stromal microenvironment was composed of tumor-associated endothelial cells, myofibroblasts with features of cancer-associated fibroblasts, and a spectrum of pericytes and vascular smooth muscle cells.

Since gene expression profiles are linked to the biological behavior of tumors, transcriptional subtypes have been defined for many tumor entities. Recently, Alcala et al. described three molecular clusters of lung neuroendocrine neoplasms based on transcriptome and methylome profiling [15]. However, it is not known to what extent information from bulk multi-omic profiling originate from tumor cells or the associated non-neoplastic immune and stromal cells. Indeed, molecular subtypes of some entities have been shown to be mainly driven by features of the tumor microenvironment, such as in colorectal cancer [25]. In our dataset, we could show that assignment of lung carcinoids to recently defined molecular clusters is not substantially driven by immune or stromal cells, but rather represent tumor-intrinsic features. Nonetheless, molecular clusters of lung neuroendocrine neoplasms have been suggested to be associated with distinct cell types of the tumor microenvironment [15]. Although the size of our dataset does not allow to define patient subgroups based on tumor microenvironment composition, we observed that the tumor of patient 1 was assigned to molecular cluster B and harbored the highest proportion of monocytes while patient 3 was assigned to cluster A1 and harbored the highest proportion of conventional dendritic cells. Exemplarily, this underlines the proposed association of molecular clusters of lung neuroendocrine neoplasms with tumor microenvironment composition [15].

Many studies have dissected the composition of the immune microenvironment of lung cancer and its potential effects on response to immune checkpoint blockade, being an important pillar in treatment of advanced disease [26]. However, the immune cellular diversity in lung carcinoids has much less been studied. It has been described that only a small proportion of carcinoids are substantially infiltrated by CD8+ T cells, which does not correlate with survival [27]. While most studies report no expression of PD-L1 in lung carcinoids at all [27, 28], some studies report a small proportion of PD-L1-postive cases and a correlation of PD-L1 expression with metastatic spread [29]. We observed that the composition of the lymphoid cell compartment closely resembled normal lung parenchyma which is in line with a recent study analyzing the lung carcinoid immune microenvironment by flow cytometry [30]. While it has been discussed that carcinoid tumors are not eligible for immune checkpoint inhibitor therapy due to their low mutational and neoantigen load [31], still, combined anti-PD1 and anti-CTL4 blockade has shown efficacy in individual advanced atypical carcinoid cases [32]. Beyond CD8+ T cells, myeloid cells in the microenvironment can modulate the response to immune checkpoint inhibitors [33]. Across all three lung carcinoids analyzed, we observed a consistent decrease in tissue-resident alveolar macrophages and an increase in monocyte-derived myeloid cells, compared to normal lung tissue, which has likewise been observed in lung adenocarcinomas [10, 14]. Monocyte-derived myeloid cells can give rise to monocyte-derived dendritic cells or monocyte-derived macrophages exhibiting different functionalities in the tumor microenvironment [20]. In our dataset, gene expression patterns favored a differentiation towards monocyte-derived macrophages. However, complementary information on protein expression is necessary to determine the lineage commitment of these cells since cell types are yet mainly defined by surface marker profiles obtained in FACS studies [20]. Furthermore, in our study, monocyte-derived myeloid cells showed low expression scores of various inflammation-related pathways, high expression of *SELENOP1*, which has been associated with M2 polarization of tumor-associated macrophages, high expression of the cytokines *CCL3* and *CCL4*, which both can exert pro- or anti-tumorigenic functions [34, 35], and high expression of all components of the C1q protein complex, which has been found to have tumor-promoting features [36, 37]. Together, we conclude that monocyte-derived myeloid cells exhibit rather non-inflammatory and pro-tumorigenic features in the lung carcinoid tumor microenvironment.

The microenvironment of neuroendocrine neoplasms often harbors a dense vascular network. We found that tumor-associated endothelial cells in lung carcinoids exhibit a distinct gene expression profile and share many highly expressed genes, such as *INSR, VWA, COL15A1, IGFBP7* and *GSN*, with tumor-associated endothelial cells of more aggressive cancers, such as lung adenocarcinoma [10, 14]. In addition, we observed a high proportion of vascular smooth muscle cells and pericytes. Antiangiogenic drugs have been in clinical trials and the VEGFR inhibitor sunitinib has been approved for therapy of pancreatic neuroendocrine tumors [31]. Interestingly, compared to normal tissues, tumor-associated pericytes expressed high levels of *RGS5*, which has been found to be overexpressed in developing pericytes during embryogenesis [38] and is associated with reduced response to VEGF inhibition in mouse models [39]. Moreover, we observed high expression of smooth muscle actin *ACTA2* in tumor-associated pericytes, which has been proposed as a marker for tumors refractory to VEGFR2 inhibition in a pancreatic neuroendocrine tumor mouse model [40]. Furthermore, the microenvironment of lung carcinoids contained a small proportion of myofibroblasts which were characterized by high TGFβ and hypoxia signaling, high expression of matrix components, matrix degrading enzymes, and marker genes such as FAP, all being features of cancer-associated fibroblasts [41, 42].

The tumor microenvironment of lung carcinoids and lung adenocarcinomas shared some features, such as tumor-associated endothelial cells, cancer-associated myofibroblasts, and monocyte-derived myeloid cells. Other microenvironmental features were specific for lung carcinoids, such as the predominance of vascular smooth muscle cells and pericytes, and the normal-like composition of the lymphoid cell compartment. Beyond these patient-overarching features of the lung carcinoid microenvironment, we observed inter-patient heterogeneity in its cellular composition. In a larger cohort of lung carcinoids profiled by bulk RNA sequencing, we found that certain cell types such as monocyte-derived myeloid cells, pericytes, and myofibroblasts might be associated with different histological subtypes of lung carcinoids (typical versus atypical) and patient prognosis. However, since the microenvironment of lung carcinoids is rather sparse yet complex, its cellular composition can only to a limited extent be inferred from bulk gene expression data. Therefore, our study forms a basis for subsequent single-cell transcriptome profiling or multiplex immunofluorescence studies of larger cohorts. In the future, a more detailed and comprehensive understanding of the tumor microenvironment could reveal specific cell types that are eligible for novel targeted therapies, and provide valuable prognostic and predictive information to improve the clinical management of lung carcinoid patients.

## Methods

### Collection of tissue specimens

Fresh tissue samples of approximately 0.1-0.5 cm^3^ of tumor tissue and normal lung parenchyma were obtained during intraoperative pathologist consultation. Informed consent was obtained from all patients were. Research was approved by vote EA4/164/19 of the ethics committee of Charité - Universitätsmedizin Berlin. Normal tissue samples of patients 4-10 have already been part of a previous study (patient 4 = P018, patient 5 = P019, patient 6 = P027, patient 7 = P029, patient 8 = P030, patient 9 = P031 and patient 10 = P033) [14].

### Tissue dissociation and single cell isolation

For transport, tissue samples were stored for max. 3 hours on ice in Tissue Storage Solution (Miltenyi). First, tissue samples were minced into pieces of max. 1 mm3 using two scalpels. Minced tissue samples were disaggregated using the Tumor Dissociation Kit, human (Miltenyi) according to the manufacturer’s protocol in a gentleMACS Octo Dissociator with heaters (Miltenyi) using the preinstalled program 37C_h_TDK_1 for 30-45 min. Subsequently, cell suspensions were filtered through 100 μm filters and kept at 4°C or on ice for all subsequent steps. Next, cells were pelleted by centrifugation at 300 g for 5 min in BSA-coated low-binding tubes, and resuspended in 1 ml ACK buffer for 60 seconds for erythrocyte lysis. Cells were washed with DMEM, again pelleted, and resuspended in PBS. After filtering the cell suspensions through 20 μm filters, debris was removed using the Debris Removal Solution (Miltenyi) according to the manufacturer’s protocol. Finally, cell concentration was determined using a Neubauer chamber.

### Single-cell RNA sequencing

Immediately after single cell isolation, 10,000 single cells per tissue sample were subjected to barcoding and library preparation, using the Chromium Single Cell 3’Reagent Kit v3 (10x Genomics) and the Chromium Controller (10x Genomics) according to the manufacturer’s protocol. Libraries were sequenced on a HiSeq 4000 Sequencer (Illumina) at average 240 mio. reads per library, resulting in average approx. 50,000 reads per cell.

### H&E and immunostaining

For hematoxylin and eosin (H&E), and immunohistochemical staining, 3-5 μm tissue sections were prepared from formalin-fixed and paraffin-embedded (FFPE) tissue.

For H&E staining, tissue sections were incubated in acidic haemalum staining solution (Waldeck) for 8 min, washed, and incubated in eosin staining solution (Sigma-Aldrich) for 2.5 min at room temperature using a Tissue-Tek Prisma Plus slide stainer (Sakura).

For antigen retrieval, tissue sections were incubated in CC2 buffer (for mouse anti-INSR) or CC1 mild buffer (for all other antibodies, Ventana Medical Systems) for 30 min at 100°C. Sections were incubated with the primary antibody for 60 min at room temperature, washed, and incubated with the secondary antibody for 30 minutes at room temperature. Antibodies were diluted in Dako Real Antibody Diluent (Dako, S2022). Staining was performed on the BenchMark XT immunostainer (Ventana Medical Systems).

The following primary antibodies were used: mouse anti-Synaptophysin (1:50, clone 27G12, Leica, NCL-L-SYNAP-299), rabbit anti-Chromogranin A (1:100, clone EP38, Epitomics, AC-0037), mouse anti-CD44 (1:50, clone DF1485, Dako, M7082), rabbit anti-EGFR (prediluted, Roche, 790-4347), rabbit anti-TFF3 (1:250, Abcam, ab108599), Rabbit anti-ERG (prediluted, clone EPR3864, Roche, 790-4576), mouse anti-Ki67 (1:50, clone MIB-1, Dako, M7240), mouse anti-INSR (1:50, clone CT-3, Invitrogen, AHR0271).

Slides were imaged using a Pannoramic SCAN 150 slide scanner (3DHISTECH).

### Single-cell gene expression analysis

#### Preprocessing

After sequencing, reads were aligned and UMIs quantified using Cellranger 3.0.2 (10x Genomics) with reference transcriptome GRCh38. All subsequent analyses were performed in R using the toolkit Seurat v4 [43], if not stated otherwise. Single-cell gene expression data of all patients were merged and filtered for the following quality parameters: 500-10,000 genes detected, 1,000-100,000 UMIs counted, fraction of mitochondrial reads <40%, and fraction of hemoglobin reads <5%. Single-cell gene expression data was normalized using the scTransform function with default parameters, and the number of UMIs per cell and the fraction of mitochondrial reads was regressed out.

#### Cell type annotation

After principal component analysis (PCA), the top 10 principal components were used for clustering and UMAP embedding of single-cell transcriptomes. Main cell types (epithelial, immune, stromal) were assigned based on cluster-wise expression of canonical cell type marker genes (resolution = 0.3, otherwise default parameters). The dataset was split into three main cell type subsets, and PCA, clustering and UMAP embedding was rerun on each subsets using the top 10 principal components and a clustering resolution of 2 with otherwise default parameters. In order to assign epithelial, immune, and stromal cell types, selected cell type marker genes according from Habermann et al. [44] and Tata et al. [45], and cell type signatures according to Vieira Braga et al. [17] and Travaglini et al. [16] were used. Clusters contaminated with epithelial or immune transcriptomes were identified by expression of *EPCAM* or *PTPRC*, respectively, and removed from the dataset prior to subsequent analyses. In the epithelial subset, cell clusters which were overrepresented in tumor tissue samples were annotated as tumor cells.

#### Differential gene expression analysis

Prior to differential gene expression analysis of epithelial cells, tumor cells from tumor samples were subset and gene expression rescaled. Immune and stromal subsets were split into lymphoid, myeloid, endothelial and fibroblastic/smooth muscle subsets, and gene expression rescaled. Next, marker genes of each cell cluster were calculated against all other clusters of the subset using the FindAllMarkers function with Wilcoxon rank-sum test and the following parameters: include only positive markers, proportion of expressing cells inside the cluster ≥ 0.25, difference between proportions of expressing cells inside and outside the cluster ≥ 0.25, log2 fold change between cells inside and outside the cluster ≥ 0.25.

#### Functional analysis

Cell cycle phases were assigned using the CellCycleScoring function. The AddModuleScore function was used to score the expression of functional relevant gene signatures: the hallmark signatures of the collection of the Broad Institute [46], naiveness, cytotoxicity and exhaustion signatures according to Guo et al. [13], and molecular cluster marker genes according to Alcala et al. [15]. For the latter, the top 50 upregulated genes in each individual molecular cluster versus the two other clusters were combined and used as gene sets. Oncogenic signaling pathway activity scores were computed using the R toolkit Progeny [47, 48] based on the top 500 genes with otherwise default parameters. The CellPhoneDB toolkit was used with default parameters to calculate potential cell-cell interactions [24]. The curated list of high-confidence ligands and receptors of oncogenic pathways can be found in ref. [14].

### Bulk gene expression and survival analysis

Bulk gene expression data was downloaded from the GitHub repository https://github.com/IARCbioinfo/DRMetrics [22] and clinical data from [15] was added. After filtering out genes located on sex and mitochondrial chromosomes, bulk gene expression data was normalized using the VarianceStabilizingTransformation function of the DESeq2 toolkit. Data on histological subtype (typical vs. atypical) was available for 75 carcinoid cases. Overall survival data was available for 76 carcinoid cases. Single-cell gene expression data was split into myeloid, lymphoid, endothelial and fibroblastic/smooth muscle subsets and rescaled. Marker genes were calculated as described above. Next, marker gene lists were used as gene sets for single-sample gene set enrichment analysis (ssGSEA) [49] of the bulk gene expression data using the gsva function of the R toolkit GSVA assuming Gaussian distribution with otherwise default parameters. For survival analyses, ssGSEA enrichment scores were dichotomized (ES > median or ≤ median). Survival curves, log-rank statistics and Cox regression were calculated using the R packages survival and survminer.

## Code and data availability

The code used for analyses is available from https://github.com/bischofp/lung_carcinoid. Gene expression count data is available from [link provided upon acceptance of the manuscript].

## Author contributions

NB, FK, PB conceived and designed the study;

PB, AT, FK, DH, AE, TD, JN, JR contributed to clinical sample acquisition and preparation;

AT, PB conducted experiments;

PB, BO, JP, DB, NB performed bioinformatic analyses;

PB, FK, NB, PJ, CF, DH, MM, CS analyzed and interpreted data and/or supervised parts of the study;

PB wrote the manuscript;

FK, NB, MM, DH revised the manuscript; all authors provided critical feedback and helped shaping the research, analysis, and manuscript.

## Acknowledgement

We thank Manuela Pacyna-Gengelbach and Barbara Meyer-Bartell for excellent technical assistance.

