## Supplemental Figures for "The single-cell transcriptional landscape of lung carcinoid tumors"

1) Charité - Universitätsmedizin Berlin, corporate member of Freie Universität Berlin and Humboldt-Universität zu Berlin, Institute of Pathology, Charitéplatz 1, 10117 Berlin, Germany
2) Berlin Institute of Health at Charité - Universitätsmedizin Berlin, Charitéplatz 1, 10117 Berlin, Germany
3) IRI Life Sciences, Humboldt University of Berlin, Philippstrasse 13, 10115 Berlin, Germany
4) Berlin Institute of Health at Charité - Universitätsmedizin Berlin, Core Unit Bioinformatics, Charitéplatz 1, 10117 Berlin, Germany
5) Institute of Pathology, LMU Munich, Thalkirchner Straße 36, 80337 München, Germany
6) Charité - Universitätsmedizin Berlin, corporate member of Freie Universität Berlin and Humboldt-Universität zu Berlin, Department of Surgery, Campus Charité Mitte and Campus Virchow-Klinikum, Charitéplatz 1, 10117 Berlin, Germany
7) Institute of Transplant Immunology, Hannover Medical School, Carl-Neuberg-Str. 1, 30625 Hannover, Germany
8) DZIF, German Center for Infectious Diseases, TTU-IICH Hannover-Braunschweig site, 38124 Braunschweig, Germany
9) German Cancer Consortium (DKTK), Partner Site Berlin, and German Cancer Research Center (DKFZ), 69120 Heidelberg, Germany
10) joint last authors

**Competing interests:** The authors declare no potential conflicts of interest.

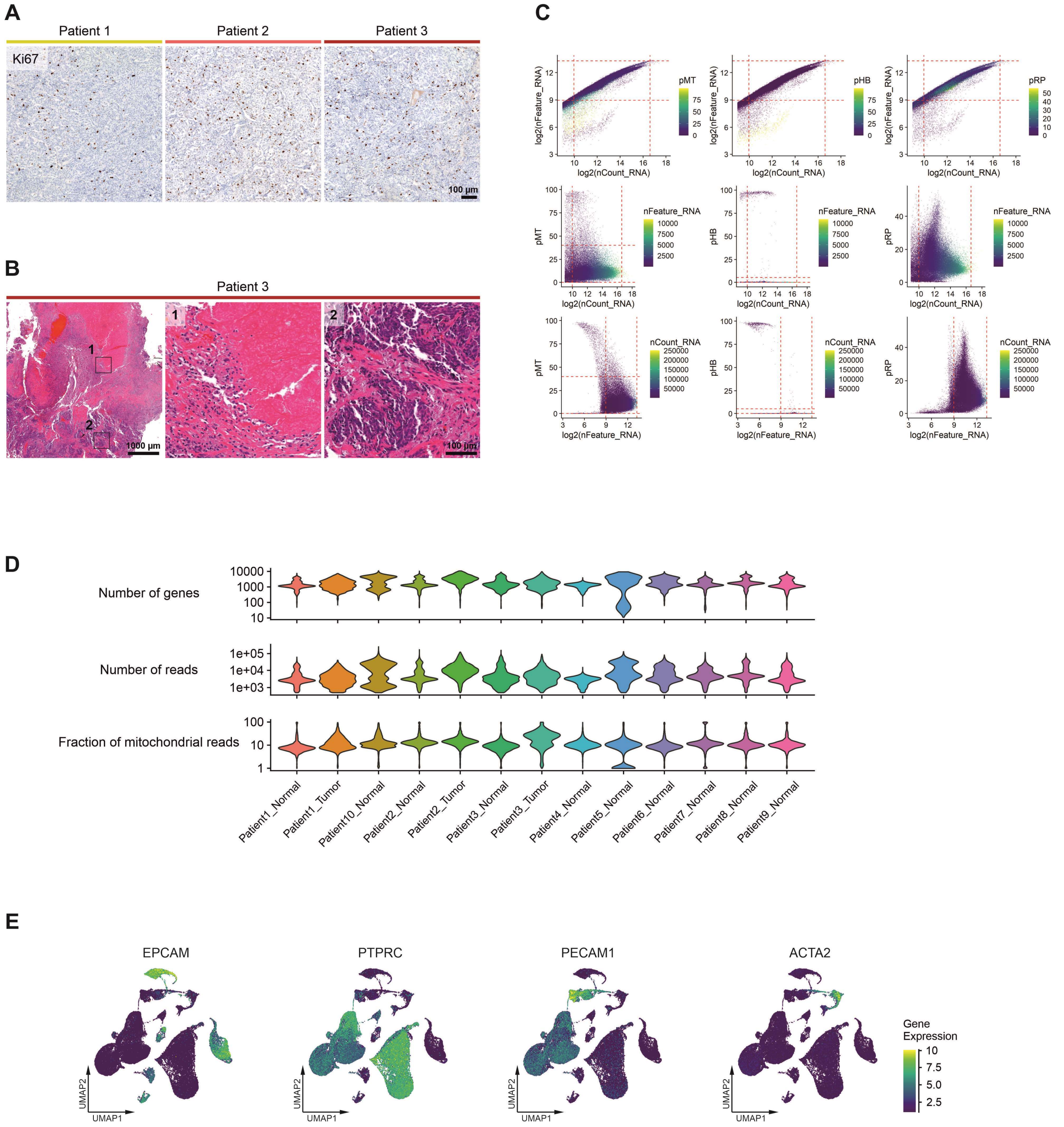


### Supplementary Figure 1

(A) Immunohistochemical staining for the proliferation marker Ki67 in all three lung carcinoid cases analyzed in the study. (B) H&E staining of the tumor necrosis in patient 3, magnified areas show (1) necrotic, and (2) vital tumor tissue. (C) Scatterplots of selected quality parameters of all single-cell transcriptomes prior to filtering, red dashed lines indicate cut-off values for filtering. (D) Violin plots of selected quality parameters of all single-cell transcriptomes prior to filtering, grouped by patient and tissue type. (E) UMAPs of all single-cell transcriptomes after filtering, colored by gene expression of selected canonical markers for epithelial (EPCAM), immune (PTPRC), and stromal cells (PECAM1, ACTA2).

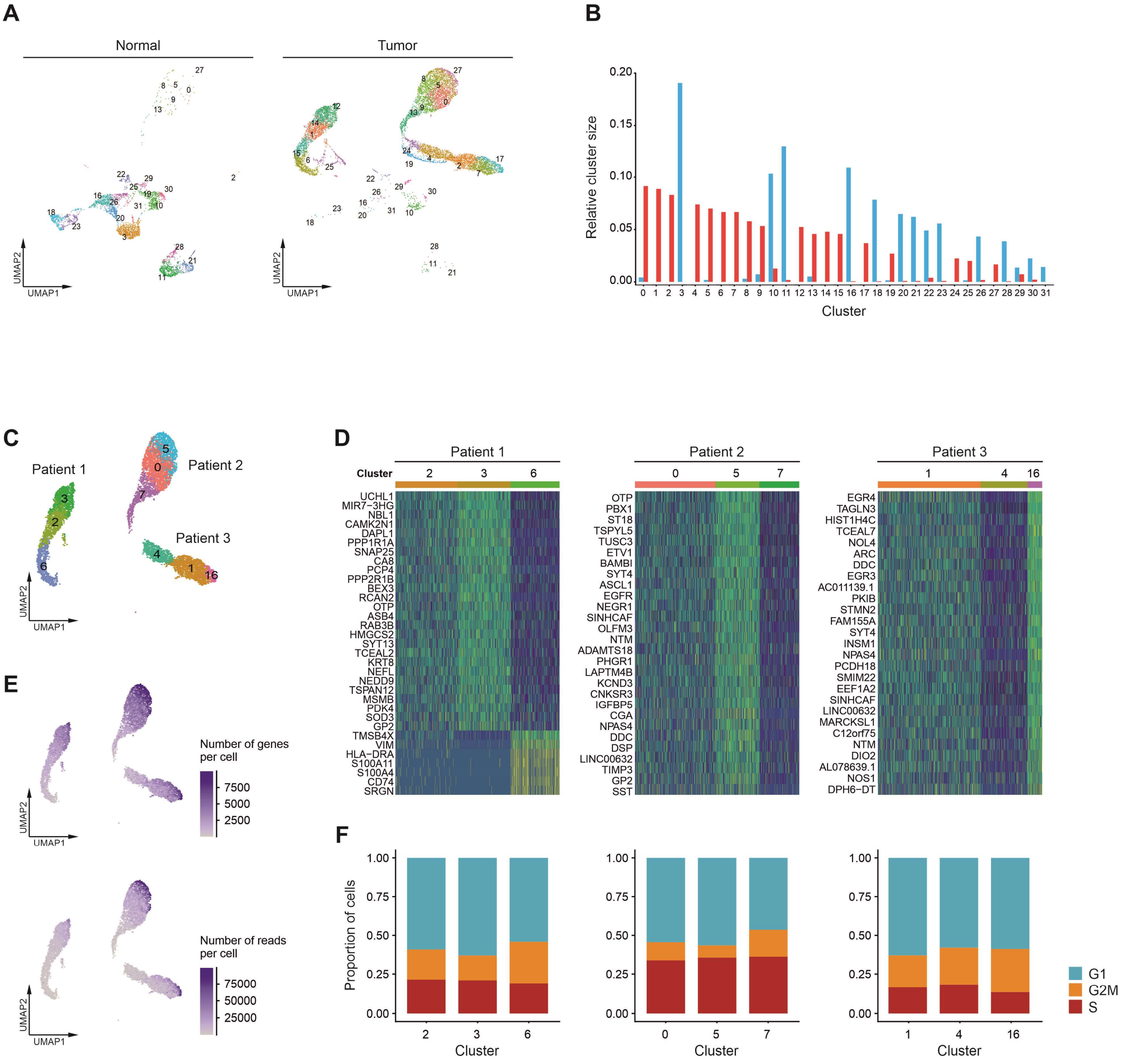


### Supplementary Figure 2

(A) UMAPs based on the top 10 principal components of all epithelial single-cell transcriptomes, split by tissue type, color-coded by cluster (resolution = 2), cluster number indicated. (B) Relative size of clusters in normal and tumor tissue samples, respectively, for clusters numbers see (A). (C-F) Epithelial cell clusters overrepresented in tumor tissue samples were assigned as tumor cell clusters and subset. (C) UMAP of tumor cell clusters, color-coded by cluster (resolution = 1), cluster number indicated. (D) Differentially expressed genes in tumor cell clusters for each patient separately, for cluster numbers see (C). (E) UMAPs of tumor cell clusters, colored by number of genes per cell and number of reads per cell. (F) Proportion of tumor epithelial cells in different cell cycle phases, grouped by clusters, for cluster numbers see (C).

A

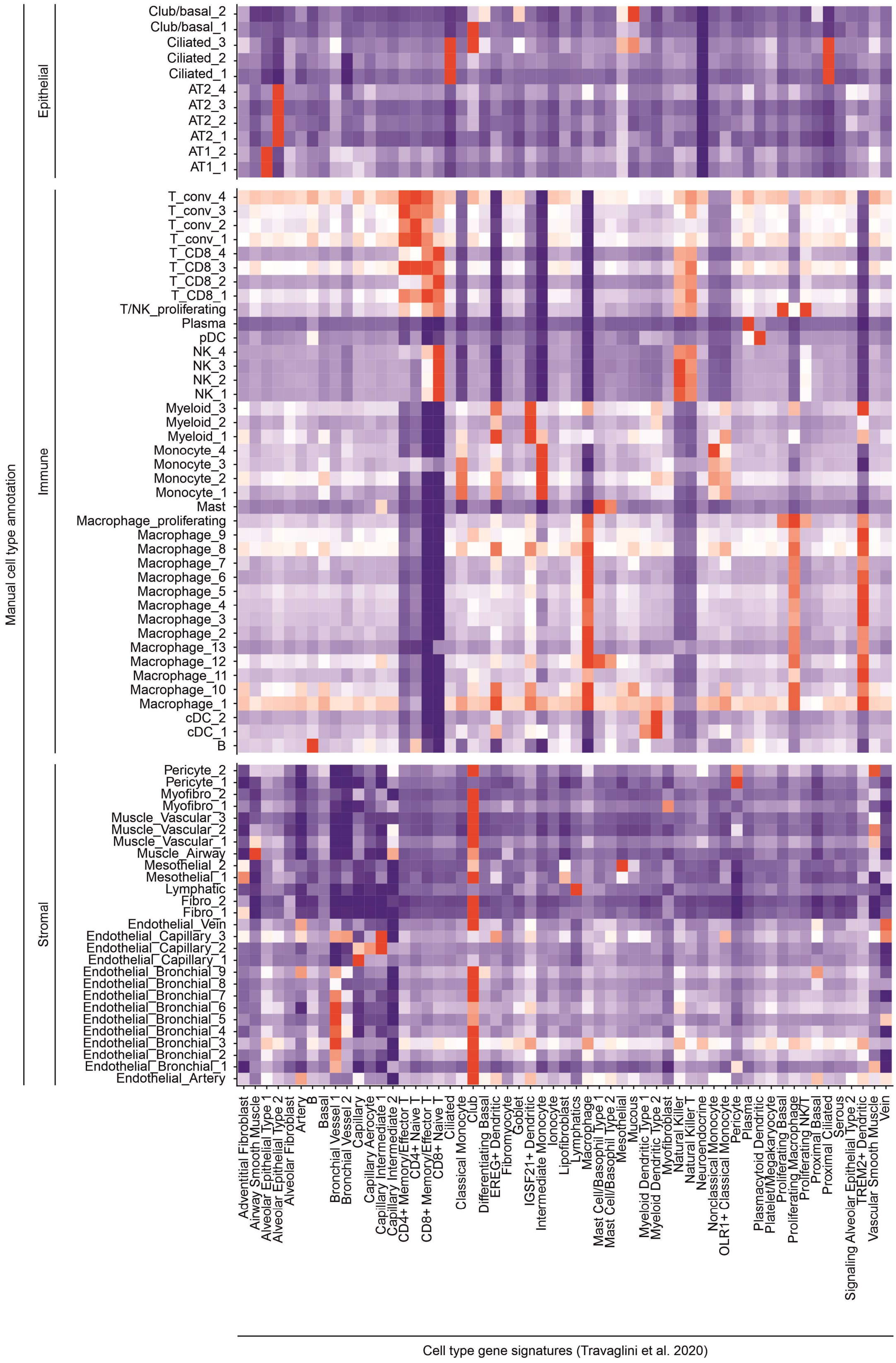
Manual cell type annotation
Epithelial
Immune
Stromal
Club/basal_2
Club/basal_1
Ciliated_3
Ciliated_2
Ciliated_1
AT2_4
AT2_3
AT2_2
AT2_1
AT1_2
AT1_1
T_conv_4
T_conv_3
T_conv_2
T_conv_1
T_CD8_4
T_CD8_3
T_CD8_2
T_CD8_1
T/NK_proliferating
Plasma
pDC
NK_4
NK_3
NK_2
NK_1
Myeloid_3
Myeloid_2
Myeloid_1
Monocyte_4
Monocyte_3
Monocyte_2
Monocyte_1
Mast
Macrophage_proliferating
Macrophage_9
Macrophage_8
Macrophage_7
Macrophage_6
Macrophage_5
Macrophage_4
Macrophage_3
Macrophage_2
Macrophage_13
Macrophage_12
Macrophage_11
Macrophage_10
Macrophage_1
cDC_2
cDC_1
B
Pericyte_2
Pericyte_1
Myofibro_2
Myofibro_1
Muscle_Vascular_3
Muscle_Vascular_2
Muscle_Vascular_1
Muscle_Airway
Mesothelial_2
Mesothelial_1
Lymphatic
Fibro_2
Fibro_1
Endothelial_Vein
Endothelial_Capillary_3
Endothelial_Capillary_2
Endothelial_Capillary_1
Endothelial_Bronchial_9
Endothelial_Bronchial_8
Endothelial_Bronchial_7
Endothelial_Bronchial_6
Endothelial_Bronchial_5
Endothelial_Bronchial_4
Endothelial_Bronchial_3
Endothelial_Bronchial_2
Endothelial_Bronchial_1
Endothelial_Artery
Adventitial Fibroblast
Airway Smooth Muscle
Alveolar Epithelial Type 1
Alveolar Epithelial Type 2
Alveolar Fibroblast
Artery
B
Basal
Bronchial Vessel 1
Bronchial Vessel 2
Capillary
Capillary Aerocyte
Capillary Intermediate 1
Capillary Intermediate 2
CD4+ Memory/Effector T
CD4+ Naive T
CD8+ Memory/Effector T
CD8+ Naive T
Ciliated
Classical Monocyte
Club
Differentiating Basal
EREG+ Dendritic
Fibromyocyte
Goblet
IGSF21+ Dendritic
Intermediate Monocyte
Ionocyte
Lipofibroblast
Lymphatics
Macrophage
Mast Cell/Basophil Type 1
Mast Cell/Basophil Type 2
Mesothelial
Mucous
Myeloid Dendritic Type 1
Myeloid Dendritic Type 2
Myofibroblast
Natural Killer
Natural Killer T
Neuroendocrine
Nonclassical Monocyte
OLR1+ Classical Monocyte
Pericyte
Plasma
Plasmacytoid Dendritic
Platelet/Megakaryocyte
Proliferating Basal
Proliferating Macrophage
Proliferating NK/T
Proximal Basal
Proximal Ciliated
Serous
Signaling Alveolar Epithelial Type 2
TREM2+ Dendritic
Vascular Smooth Muscle
Vein
Cell type gene signatures (Travaglini et al. 2020)

B

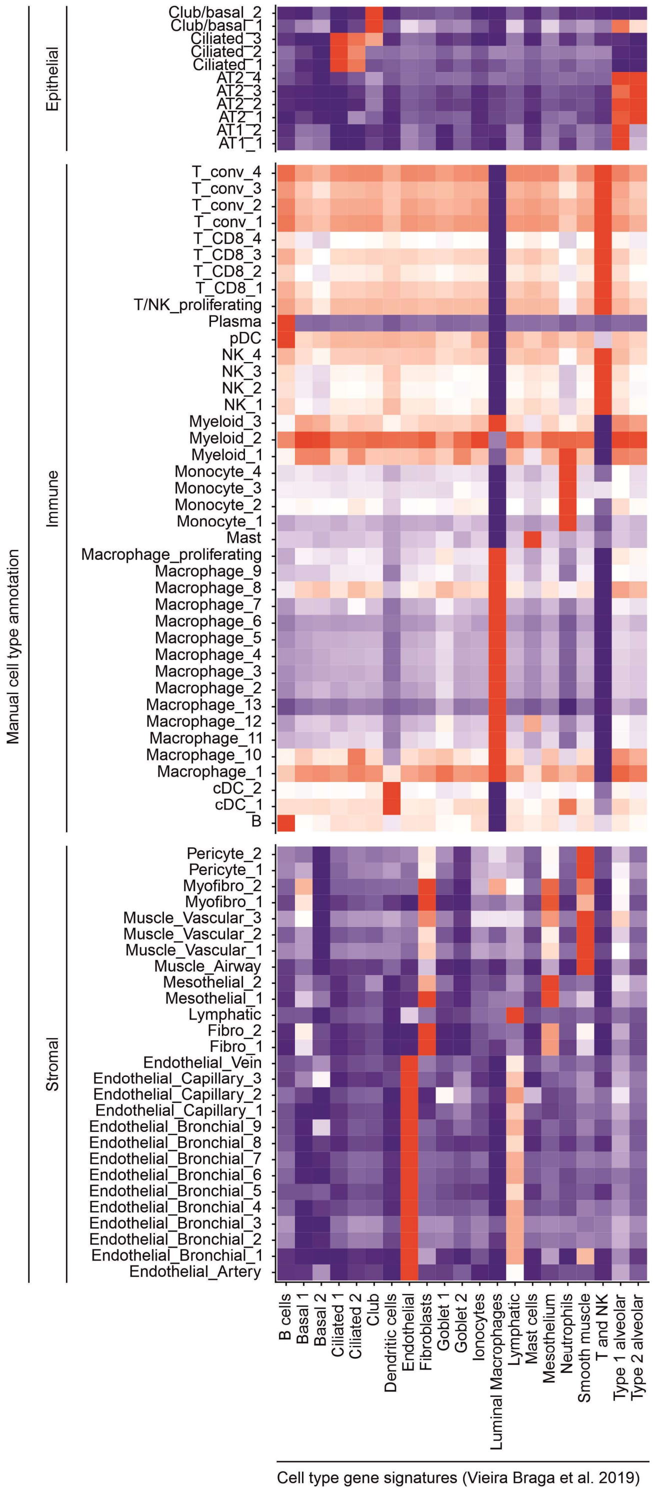


**Supplementary Figure 3**

(A+B) Mean module scores of cell type gene signatures according to (A) Travaglini et al. and (B) Vieira Braga et al. in epithelial, immune and stromal cell clusters as annotated manually.

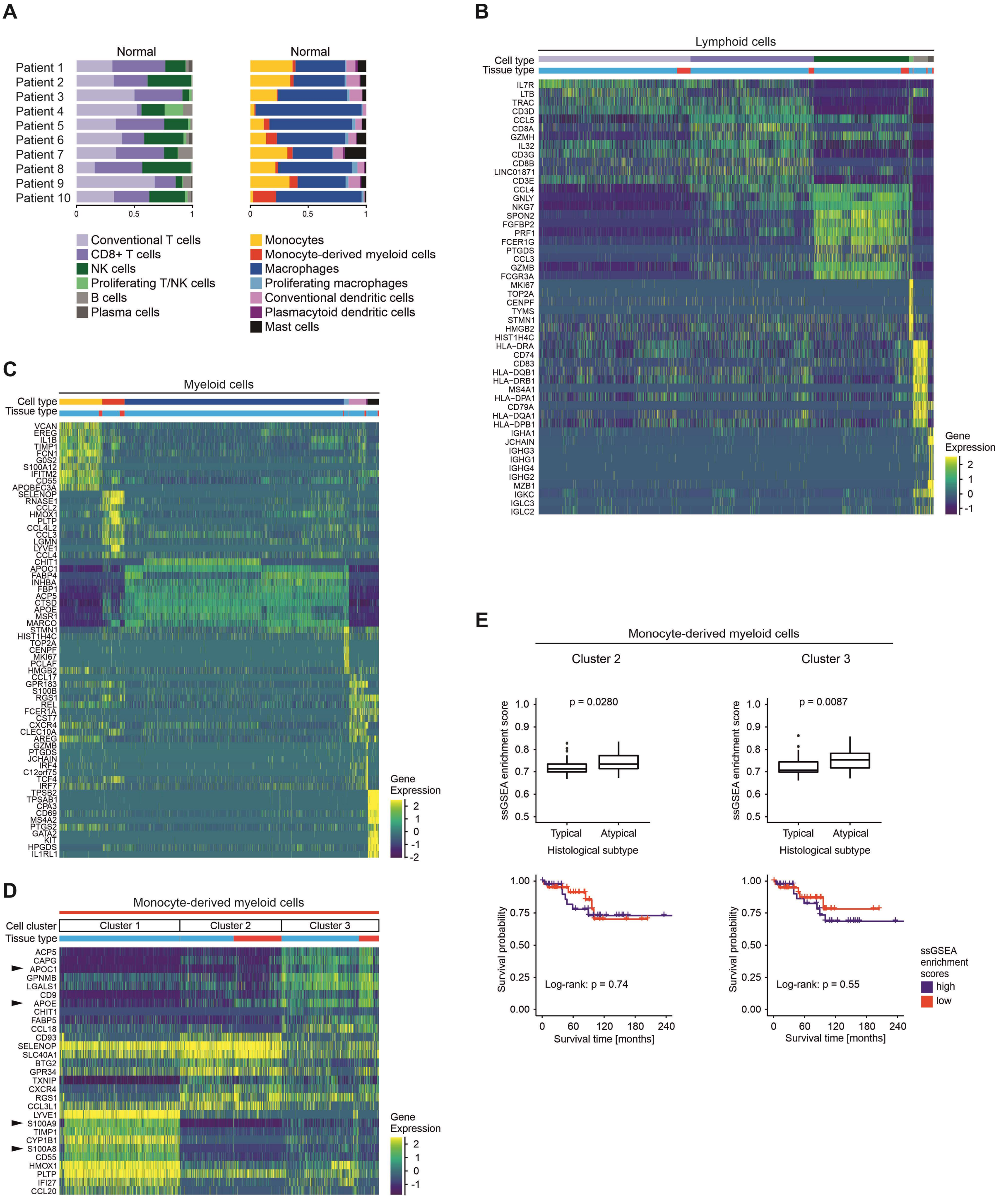


### Supplementary Figure 4

(A) Proportions of lymphoid and myeloid cell types per patient in normal tissue samples. (B+C) Differentially expressed genes of (B) lymphoid and (C) myeloid cell types, top 10 genes showed per cell type. (D) Differentially expressed genes of monocyte-derived myeloid cell clusters, top 10 genes showed per cell cluster, black arrowheads indicate genes mentioned in the main text. (E) ssGSEA enrichment scores in 75 lung carcinoids analyzed by bulk gene expression profiling grouped by histological subtype, two-sided t-test statistics. Kaplan-Meier overall survival curves of 76 lung carcinoid patients grouped by ssGSEA enrichment scores, log-rank statistics.

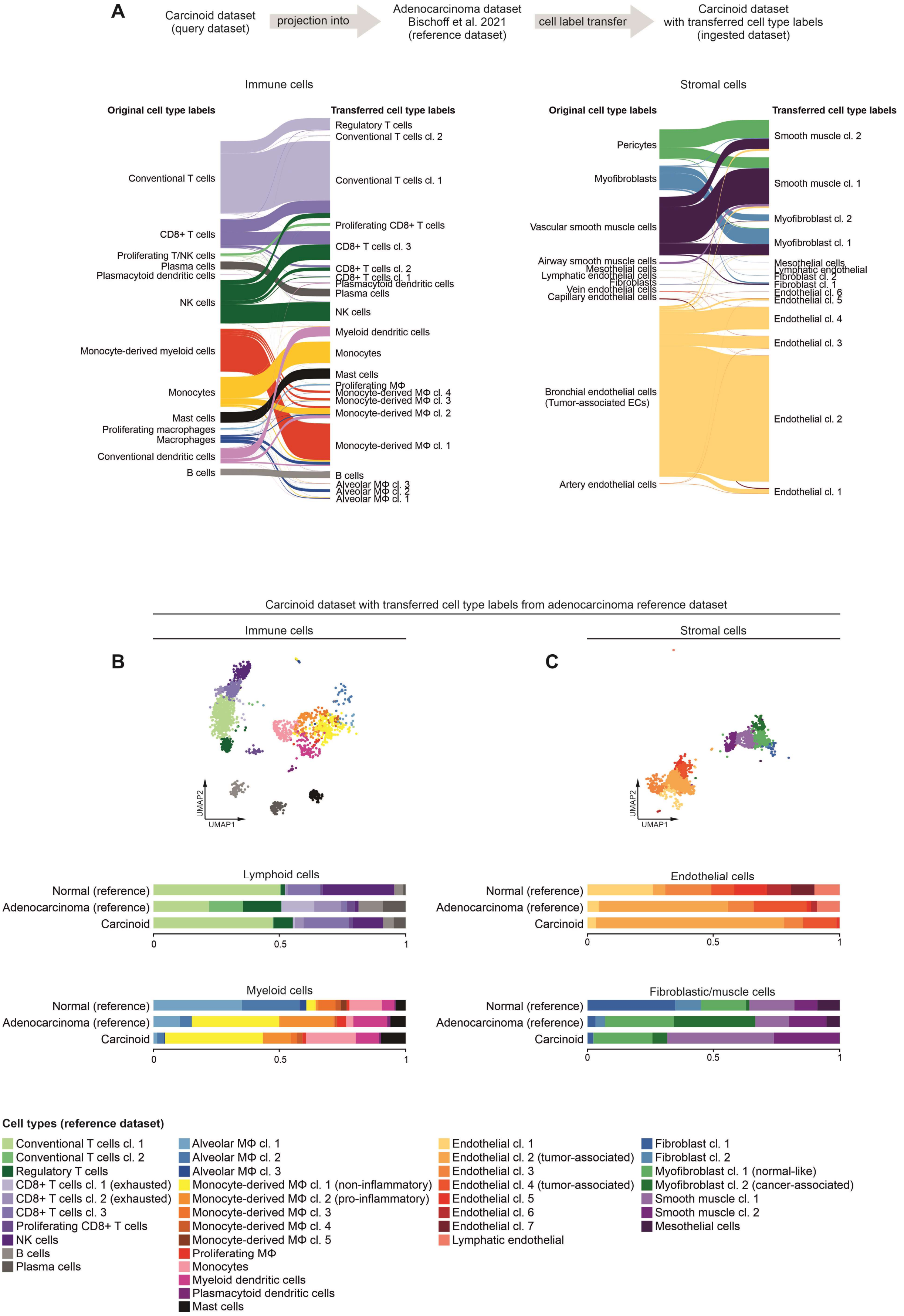
A
Carcinoid dataset
(query dataset)
projection into
Adenocarcinoma dataset
Bischoff et al. 2021
(reference dataset)
cell label transfer
Carcinoid dataset
with transferred cell type labels
(ingested dataset)
Immune cells
Original cell type labels
Transferred cell type labels
Conventional T cells
CD8+ T cells
Proliferating T/NK cells
Plasma cells
Plasmacytoid dendritic cells
NK cells
Monocyte-derived myeloid cells
Monocytes
Mast cells
Proliferating macrophages
Macrophages
Conventional dendritic cells
B cells
Regulatory T cells
Conventional T cells cl. 2
Conventional T cells cl. 1
Proliferating CD8+ T cells
CD8+ T cells cl. 3
CD8+ T cells cl. 2
CD8+ T cells cl. 1
Plasmacytoid dendritic cells
Plasma cells
NK cells
Myeloid dendritic cells
Monocytes
Mast cells
Proliferating MΦ
Monocyte-derived MΦ cl. 4
Monocyte-derived MΦ cl. 3
Monocyte-derived MΦ cl. 2
Monocyte-derived MΦ cl. 1
B cells
Alveolar MΦ cl. 3
Alveolar MΦ cl. 2
Alveolar MΦ cl. 1
Stromal cells
Original cell type labels
Transferred cell type labels
Pericytes
Myofibroblasts
Vascular smooth muscle cells
Airway smooth muscle cells
Mesothelial cells
Lymphatic endothelial cells
Fibroblasts
Vein endothelial cells
Capillary endothelial cells
Bronchial endothelial cells
(Tumor-associated ECs)
Artery endothelial cells
Smooth muscle cl. 2
Smooth muscle cl. 1
Myofibroblast cl. 2
Myofibroblast cl. 1
Mesothelial cells
Lymphatic endothelial
Fibroblast cl. 2
Fibroblast cl. 1
Endothelial cl. 6
Endothelial cl. 5
Endothelial cl. 4
Endothelial cl. 3
Endothelial cl. 2
Endothelial cl. 1
Carcinoid dataset with transferred cell type labels from adenocarcinoma reference dataset
Immune cells
Stromal cells
B
C
UMAP2
UMAP1
UMAP2
UMAP1
Lymphoid cells
Normal (reference)
Adenocarcinoma (reference)
Carcinoid
0
0.5
1
Endothelial cells
Normal (reference)
Adenocarcinoma (reference)
Carcinoid
0
0.5
1
Myeloid cells
Normal (reference)
Adenocarcinoma (reference)
Carcinoid
0
0.5
1
Fibroblastic/muscle cells
Normal (reference)
Adenocarcinoma (reference)
Carcinoid
0
0.5
1
Cell types (reference dataset)
Conventional T cells cl. 1
Conventional T cells cl. 2
Regulatory T cells
CD8+ T cells cl. 1 (exhausted)
CD8+ T cells cl. 2 (exhausted)
CD8+ T cells cl. 3
Proliferating CD8+ T cells
NK cells
B cells
Plasma cells
Alveolar MΦ cl. 1
Alveolar MΦ cl. 2
Alveolar MΦ cl. 3
Monocyte-derived MΦ cl. 1 (non-inflammatory)
Monocyte-derived MΦ cl. 2 (pro-inflammatory)
Monocyte-derived MΦ cl. 3
Monocyte-derived MΦ cl. 4
Monocyte-derived MΦ cl. 5
Proliferating MΦ
Monocytes
Myeloid dendritic cells
Plasmacytoid dendritic cells
Mast cells
Endothelial cl. 1
Endothelial cl. 2 (tumor-associated)
Endothelial cl. 3
Endothelial cl. 4 (tumor-associated)
Endothelial cl. 5
Endothelial cl. 6
Endothelial cl. 7
Lymphatic endothelial
Fibroblast cl. 1
Fibroblast cl. 2
Myofibroblast cl. 1 (normal-like)
Myofibroblast cl. 2 (cancer-associated)
Smooth muscle cl. 1
Smooth muscle cl. 2
Mesothelial cells

**Supplementary Figure 5**

(A) Lung carcinoid single-cell gene expression data was projected into a lung adenocarcinoma reference dataset (https://doi.org/10.1038/s41388-021-02054-3) using the “ingest” function of the Scanpy toolkit to transfer cell type labels and UMAP coordinates from the reference dataset to the lung carcinoid dataset. (B-C) UMAPs and proportion of cell types of (B) immune and (C) stromal single-cell transcriptomes of the lung carcinoid dataset, UMAP coordinates computed from lung adenocarcinoma reference dataset, color-coded by cell type as transferred from lung adenocarcinoma reference dataset. Functional annotation of selected cell clusters indicated in brackets, for further characterization of cell clusters, see https://doi.org/10.1038/s41388-021-02054-3.

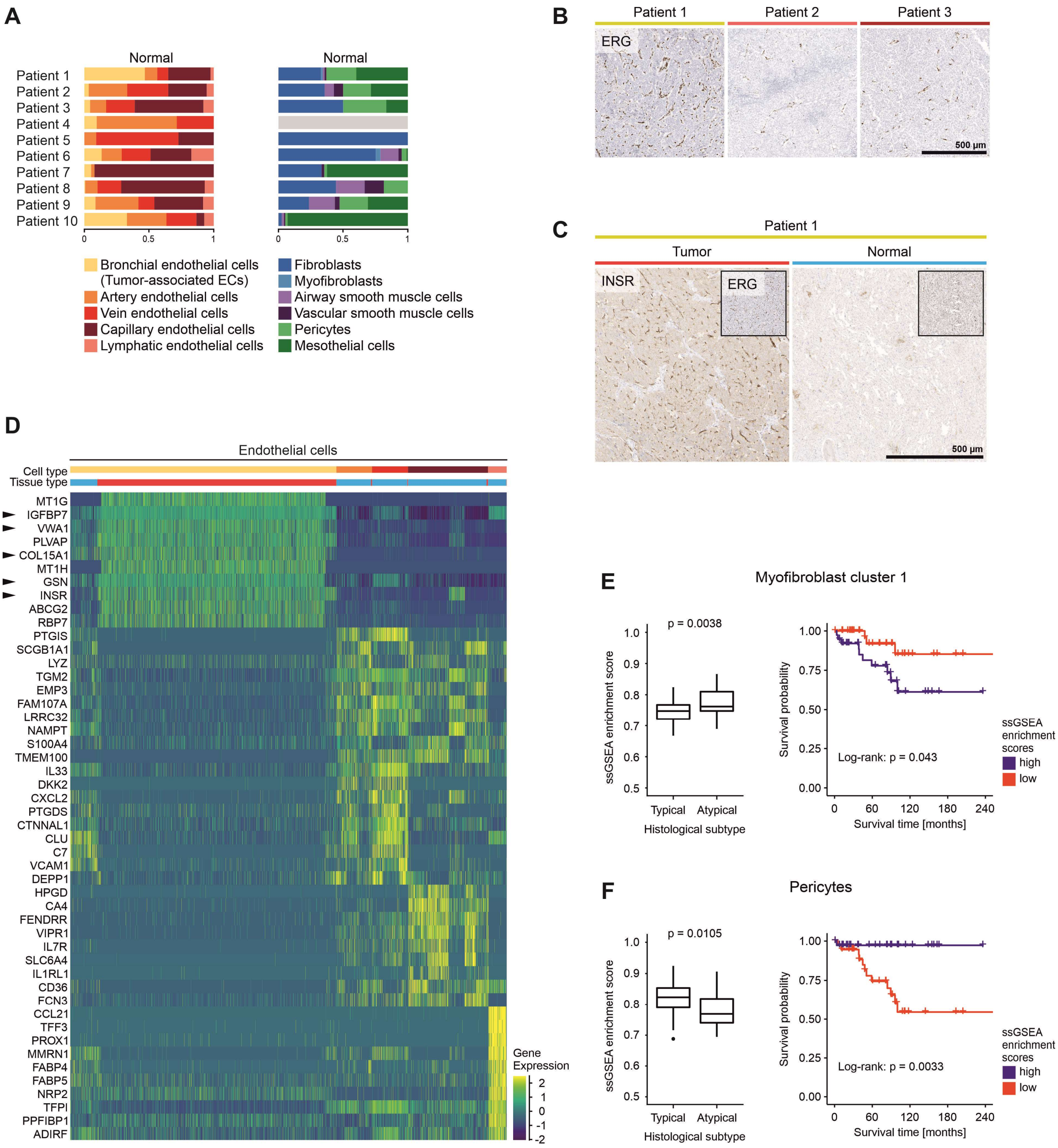


**Supplementary Figure 6**

(A) Proportions of endothelial and fibroblastic/smooth muscle cell types per patient in normal tissue samples. (B+C) Immunohistochemical staining for (B) the endothelial marker ERG and (C) the tumor-associated endothelial marker INSR. (D) Differentially expressed genes of endothelial cell types, top 10 genes showed per cell type, black arrowheads indicate genes mentioned in the main text. (E+F) ssGSEA enrichment scores of (E) marker genes of myofibroblast cluster 1 and (F) pericyte marker genes in 75 lung carcinoids analyzed by bulk gene expression profiling grouped by histological subtype, two-sided t-test statistics. Kaplan-Meier overall survival curves of 76 lung carcinoid patients grouped by ssGSEA enrichment scores, log-rank statistics.

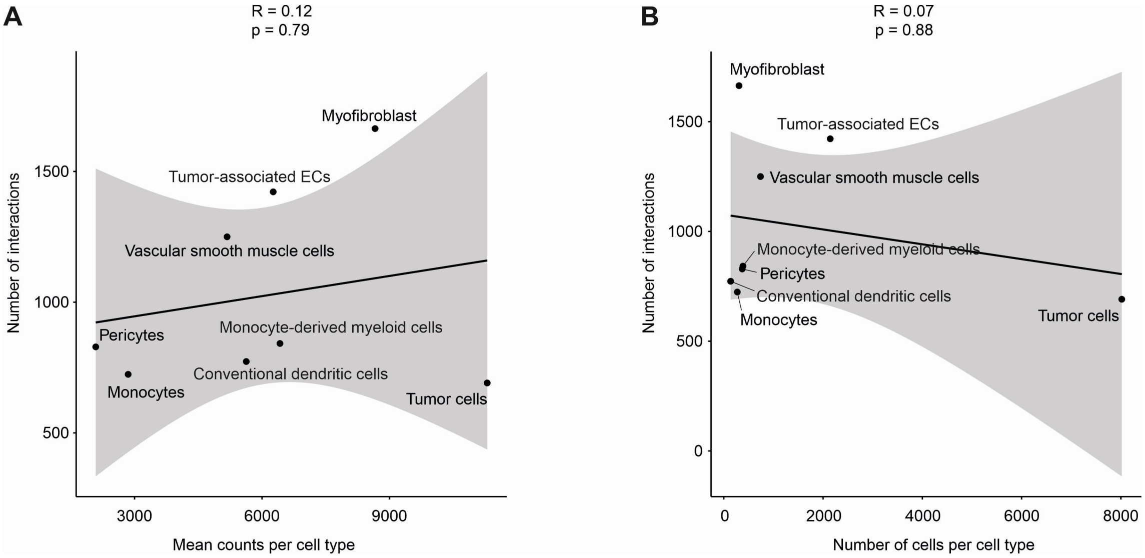


**Supplementary Figure 7**

(A+B) Correlation of the number of potential cell-cell interactions, computed using the CellPhoneDB algorithm, with (A) the mean number of mRNA counts per cell type, and with (B) the total number of cells per cell type, Pearson's correlation statistics.
